# The integrated stress response pathway activates 4E-BP to bias mRNA translation and boost antimicrobial peptide synthesis in response to bacterial infection

**DOI:** 10.1101/147629

**Authors:** Deepika Vasudevan, Nicholas K. Clark, Jessica Sam, Beatrix Ueberheide, Michael T. Marr, Hyung Don Ryoo

## Abstract

Pathogenic bacterial infection imposes considerable cellular stress on the host and often leads to attenuation of mRNA translation. In this translation-suppressive environment, it is unclear how the host synthesizes various antimicrobial peptides (AMPs) to mount innate immune response. Here, we use *Drosophila* as a model to demonstrate that AMP production during infection relies on a translation bias mechanism mediated by the inhibitor of cap-dependent translation 4E-BP (*Drosophila* Thor), and the AMP 5’UTRs that can undergo cap-independent translation. We found that 4E-BP is induced upon infection with the pathogenic bacteria *Ecc15* by the stress-responsive transcription factor ATF4, and its upstream kinase GCN2. Moreover, loss of *gcn2, atf4* or *4e-bp* compromised immunity against *Ecc15.* In 4E-BP mutants, the transcriptional induction of AMPs after infection was unaffected, while the protein levels of AMPs were substantially reduced in their hemolymph. Analysis of the 5’UTRs of AMPs using cell-based bicistronic reporters and *in vitro* translation analysis indicated that AMPs are translated in a cap-independent mechanism. Analysis of bicistronic reporters in the presence of 4E-BP indicate that infection enhances cap-independent translational activity associated with AMP 5’UTRs, accounting for enhanced AMP translation during infection.

**Highlights:**

- 4E-BP is transcriptionally induced by GCN2/ATF signaling in response to bacterial infection
- 4E-BP mutants show unaltered antimicrobial peptide (AMP) transcript levels, but have reduced AMP translation
- AMP 5’UTRs are translated cap-independently
- Translation bias by 4E-BP drives cap-independent AMP translation

## Introduction

Various types of stresses dampen protein synthesis by reducing the availability of critical components of translation initiation factors. One such example is mediated by stress-activated kinases that phospho-inactivate eIF2α, which is one of three subunits of the eIF2 complex whose normal role is to bring initiator methioninyl tRNA (Met-tRNAi^Met^) to the 43S ribosomal subunit [1]. Even under such inhibitory conditions, a subset of transcripts containing overlapping upstream ORFs (uORFs) in their 5’UTR, such as ATF4 (*cryptocephal* or *crc* in *Drosophila)*, is favorably synthesized. ATF4 responds to cellular stress by transcriptionally inducing various stress responsive transcripts. As metazoans have multiple eIF2α kinases that active this pathway, it is often referred to as the ‘integrated stress response’ (ISR)[2, 3].

Translational inhibition mechanisms associated with viral infection were first observed over fifty years ago in ascites-tumor cells infected with encephalomyocarditis virus [4] and since then this phenomenon has been extended to most viral infections. Such translational inhibition is mediated in part by PKR, one of the four eIF2α kinases in mammals that is activated by dsRNAs[5]. While not as well recognized, literature reports that pathogenic bacterial infection also causes translation inhibition in various infection models from *C. elegans* to mammals (reviewed comprehensively in [6]). Examples include a report implicating GCN2, an eIF2α kinase that responds to amino acid deprivation, in triggering an mRNA translational block of the host *Drosophila* infected with *Pseudomonas entomophila* (*P.e)* [7]. Other studies have reported the activation of the cap-dependent translational inhibitor 4E-BP (*Thor* in *Drosophila)* in response to bacterial pathogens [7–9]. Inhibition by 4E-BP impinges at the 5’-cap of eukaryotic mRNAs, where the 7-methylguanosine moiety is recognized by eIF4E (as part of the eIF4F complex) to recruit other translation initiation factors, and eventually the 43S ribosomal subunit. 4E-BP, or eIF<u>4E</u>-<u>b</u>inding <u>p</u>rotein, imposes translation inhibition by binding to <u>e</u>ukaryotic <u>i</u>nitiation factor <u>4E</u> (eIF4E), thereby specifically inhibiting cap-dependent translation [10]. As 4E-BP mutants are immune compromised [9], it is thought that induction of 4E-BP in response to bacterial infection is a host adaptation mechanism [8]. Such hypothesis that translation inhibition is a host defense mechanism raises important questions: how does translational inhibition aid the host mount an immune response that relies to on antimicrobial peptide (AMP) expression? And what signaling pathways mediate *4E-BP* induction in response to bacterial infection? Here, we identify ISR pathway as the mediator of 4E-BP induction in *Drosophila* during infection. We also show how 4E-BP paradoxically stimulates the synthesis AMPs as part of the innate immune response to bacterial infection in *Drosophila*. In a broader sense, we provide an *in vivo* demonstration of a mechanism that explains the long-standing issue of how eukaryotic cells manage their stress responsive gene expression programs under conditions of translational inhibition.

## Results

### 4E-BP is transcriptionally induced by ATF4 during infection in the fat body and gut

Independent studies have established that bacterial infection blocks the host *Drosophila* translational machinery through GCN2 [7] and 4E-BP activation [8, 11]. 4E-BP transcription is induced during this process, but the mediating signaling pathway responsible for this induction remains unknown. Recently, we and others recently reported that the eIF2α-kinase responsive transcription factor, ATF4, regulates 4E-BP transcription via its intronic element in *Drosophila* [12, 13], which prompted us to examine the possible role of ISR in 4E-BP induction. This regulation was exploited to generate the 4E-BP^intron^-dsRed reporter, which allows visualization of ATF4 activity in live *Drosophila* tissues [13]. We used this reporter to test if ATF4 is active in the context of infection, by enterically infecting 3^rd^ instar larvae with the non-lethal gram-negative bacterial pathogen, *Ecc15*. In response to infection, dsRed expression was elevated in the larval gut (Fig. 1A’, B’) and also in the fat body (Fig. 1C’, D’), which is known to be an auxiliary immune-response tissue. As reported previously [8], we saw 4E-BP transcript induction in response to *Ecc15* infection and this induction was suppressed in the background of the homozygous ATF4 hypomorphic mutant, *crc^1^* [14] (Fig. 1E) further corroborating that ATF4 mediates 4E-BP induction during infection. 4E-BP induction was also suppressed in mutants for another 4E-BP transcription factor, FOXO [15], (Fig. 1E) suggesting that multiple transcription factors may regulate 4E-BP induction during infection.

**Fig. 1:**
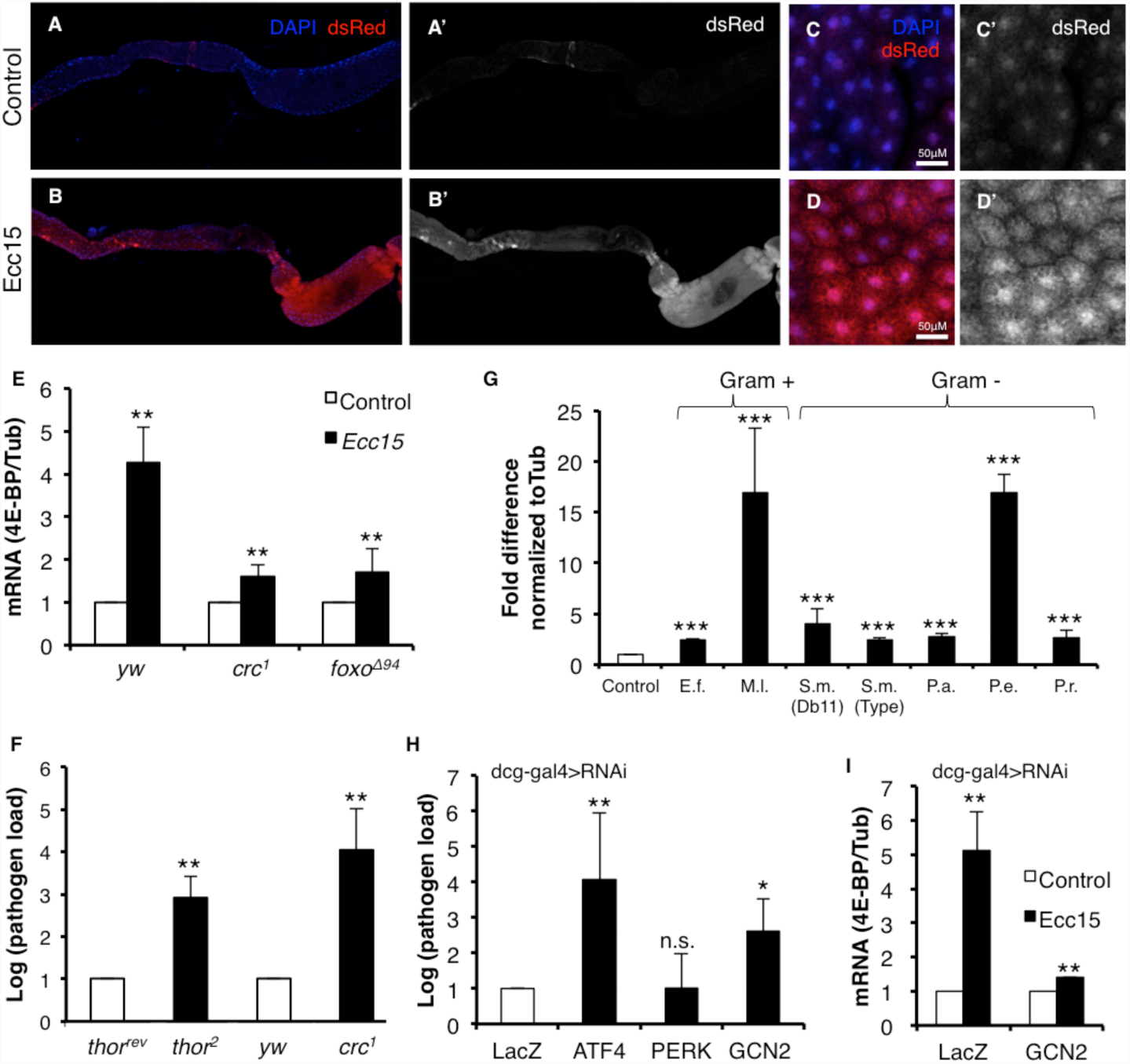
GCN2-ATF4 signaling induces 4E-BP during infection. 1A-D. 4E-BP^intron^-dsRed expression in the uninfected control larva (A, A’, C, C’) and in larva infected with *Ecc15* (B, B’, D, D’). Shown are the gut (A, A’, B, B’) and the fat body (C, C’, D, D’). A’, B’, C’, D’ are dsRed channel only images. 1E.qPCR analysis of 4E-BP expression from control larvae (genotype: *yw*), ATF4 mutants (*crc^1^* homozygotes) and FOXO mutants (*foxo^Δ94^* homozygotes) infected with *Ecc15*. 1F. Systemic pathogen load assay measuring *Ecc15* levels in 4E-BP mutants (*thor^2^* homozygotes) with 4E-BP revertants (*thor^rev^*) as control, and in homozygotic *crc^1^* with *yw* as control. 1G.qPCR analysis of 4E-BP induction in *yw* control larvae infected with various gram positive and gram negative pathogens. (*E.f.* = *Enterococcus faecalis, M.l.* = *Micrococcus luteus, S.m.* = *Serratia marcescens, P.a.* = *Pseudomonas aeruginosa, P.e.* = *Pseudomonas entomophila, P.r.* = *Providencia rettgrei*).1H. Systemic pathogen load assay on larvae expressing *uas-LacZ^RNAi^* (negative control), *-ATF4^RNAi^*, *-PERK^RNAi^*, -*GCN2^RNAi^* driven by a fat body specific driver (*dcg-gal4*).1I. qPCR analysis of 4E-BP induction in larvae expressing *uas-LacZ^RNAi^* (negative control) or -*GCN2^RNAi^* driven by a fat body specific driver (*dcg-gal4*). All data are mean of at least 3 independent experiments, error bars indicate standard error. *=p<0.05, **=p<.01, ***=p<0.001. All p-values are calculated w.r.t. respective controls.

If ATF4 was indeed upstream of 4E-BP in the context of infection, we would anticipate ATF4 mutants to be immune-compromised similar to the reported phenotype of the 4E-BP null mutant *thor^2^* [8]. We tested this using systemic pathogen load assays to measure *Ecc15* levels in the larvae after infection. *crc^1^* homozygotic mutants had higher systemic pathogen load, and thus were immune-compromised similar to 4E-BP mutants (Fig. 1F). While *Ecc15* is a reliable *Drosophila* pathogen, we further examined if the induction of 4E-BP occurred in response to other pathogens, including gram-positive bacterial (*Enterococcus faecalis*, *Micrococcus luteus*) and other gram-negative bacteria (*Serratia marcescens*, *Pseudomonas aeruginosa*, *Pseudomonas entomophila*, *Providencia rettgrei*) (Fig. 1G). We found 4E-BP induction with all pathogens we tested, albeit to different extents, suggesting that ATF4/4E-BP axis activation occurs as part of a general immune response to pathogenic bacteria.

### GCN2 signaling activates ATF4-mediated 4E-BP induction in the fat body

ATF4 can be activated by either of two known eIF2α-kinases in *Drosophila*: the ER stress responsive PERK or the amino acid deprivation activated kinase GCN2[16]. We sought to examine which of these two kinases lies upstream of ATF4 during infection. Upon further examination of the transcriptional induction of 4E-BP post-infection, we see increased 4E-BP transcripts both in the fat body and in the gut (Fig. S1). Fat body is the primary site of AMP synthesis in response to infection [17, 18], and fat body specific knockdown of GCN2 with the *dcg-gal4* driver, but not of PERK, resulted in an increased systemic pathogen load upon *Ecc15* infection (Fig. 1H). By extension, knockdown of GCN2 also resulted in the loss of 4E-BP induction in response to infection (Fig. 1I).

### Levels of AMP protein, but transcripts, are reduced in 4E-BP mutants

To gain insight into why 4E-BP mutants were immune-compromised, we asked whether 4E-BP regulated the canonical innate immune response pathways. In response to infection, various Toll and immune deficiency pathway receptors are activated and initiate a signaling cascade which culminates in the transcriptional induction of AMPs [19]. AMPs are required for combating pathogen load with different classes of AMPs targeting different aspects of pathogen physiology. We observed that the transcriptional induction of AMPs such as Drosomycin, Diptericin and Attacins is unaffected in *thor^2^* homozygotic mutants (Fig. 2A). However, western blotting of hemolymph collected from infected and uninfected *thor^2^* homozygotic larvae showed a significant reduction in the protein levels of Attacins (Fig. 2B). Further mass spectrometric analysis of hemolymph from these larvae indicated a reduction in the levels of other AMPs such as Drosocin, Diptericin B and Metchnikowin (Fig. 2C) when normalized to larval serum protein. Together, these data suggested that 4E-BP mutants were immune-compromised because of reduced AMP synthesis.

**Fig. 2:**
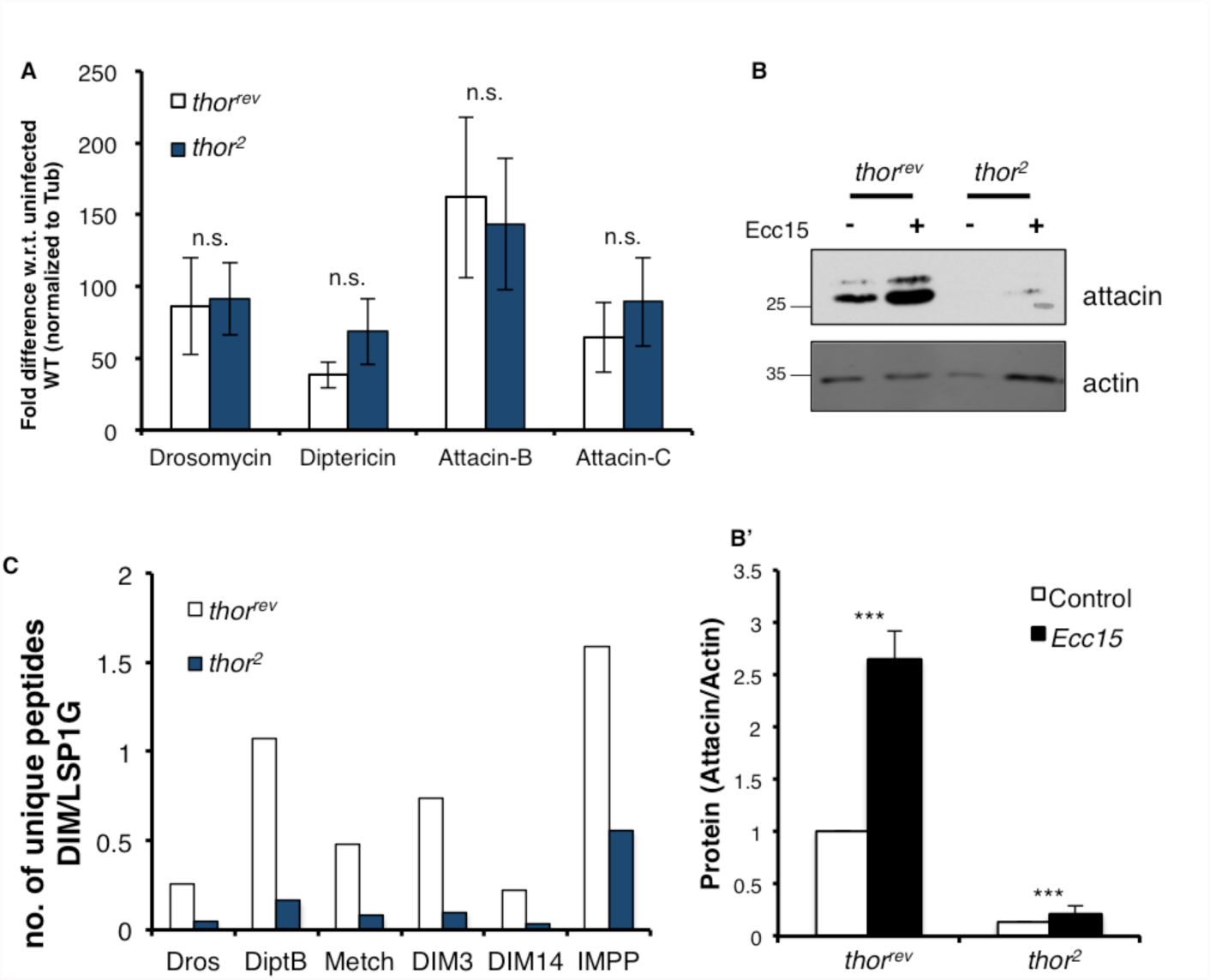
Levels of AMP protein, but not transcripts, are negatively affected in 4E-BP mutants. 2A. qPCR analysis of AMP induction in homozygotic *thor^rev^*(white) and *thor^2^* (blue) infected with *Ecc15*. The AMPs tested are represented along the X-axis. Data were normalized to the uninfected control of the respective genotype. 2B. Western blot analysis of hemolymph collected from homozygotic *thor^rev^* and *thor^2^* infected with *Ecc15* with Attacin A antibody (top panel) and a loading control, actin (bottom panel). Quantification of the blot is shown in graph below with all values normalized to *thor^rev^* control. 2B’. Quantiation of 2B. 2C. Mass spectrometry analysis of fractionated hemolymph collected from *thor^rev^*(white) and *thor^2^*(blue) infected with *Ecc15*. Data represent the number of unique peptides observed for each AMP or DIM shown on the X-axis as normalized to peptides from larval serum protein 1G (LSP1G). (Dros = Drosocin, DiptB = Diptericin B, Metch = Metchnikowin, DIM = Drosophila immune molecule, IMPP = Immune induced peptide precursor). Also see Fig. S2. Where applicable, data are mean of at least 3 independent experiments, error bars indicate standard error. All data are mean of at least 3 independent experiments, error bars indicate standard error. *=p<0.05, **=p<.01, ***=p<0.001. All p-values are calculated w.r.t. respective controls.

### 5’UTRs of Drosomycin and Attacin A have cap-independent translation activity

Loss of 4E-BP is expected to enhance cap-dependent translation, but our data suggests that AMP synthesis is somehow reduced under these conditions. This observation prompted us to examine the possibility that AMPs are translated not through eIF4E-mediated translation, but through other unconventional mechanisms. There are several known mechanisms of translation that can bypass eIF4E requirement. One of those is through the presence of an internal ribosomal entry site (IRES) element in the 5’UTR of mRNAs [20], which would allow those transcripts to recruit the 43s preinitiation complex independent of eIF4E and thus can bypass translational regulation by 4E-BP. To test the 5’UTRs of the AMPs for possible IRES elements we performed bicistronic assays in *Drosophila* S2 cells where cap-dependent translation is reported by Renilla luciferase and IRES-mediated translation is reported by Firefly luciferase (schematic, Fig. 3A). Luciferase activity measured in S2 cells expressing these bicistronic reporter constructs show that 5’UTRs of Drosomycin and Attacin A when inserted between the two cistrons in the forward orientation allow the translation of the Firefly luciferase (Fig. 3A). Interestingly, we also found that the 5’UTR of 4E-BP itself also scored positively in the bicistronic assay (Fig. 3A). Surprisingly, the cap-independent activities of these 5’UTRs were higher than that of Hepatitis C virus (HCV) IRES [21] used as a positive control. To further validate the 5’ cap-requirement of the AMP transcripts we performed *in vitro* cap competition assays by measuring translation of 5’UTR-Firefly luciferase reporters in rabbit reticulocyte lysates (RRL). While control transcripts are negatively affected by the addition of excess cap compound (m^7^G) that compete for cap-dependent translational machinery, transcripts containing the 5’UTR of Drosomycin, Attacin and 4E-BP were indifferent to the presence of the excess m^7^G (Fig. 3B). While cap-dependent transcripts would benefit from the removal of other competing mRNA from the RRL, translation of AMP 5’UTR reporters were indifferent to removal of competing mRNAs using micrococcal nuclease from RRL (Fig. 3C). Together these data support the idea that certain AMP transcripts can be translated independent of eIF4E.

**Fig. 3.**
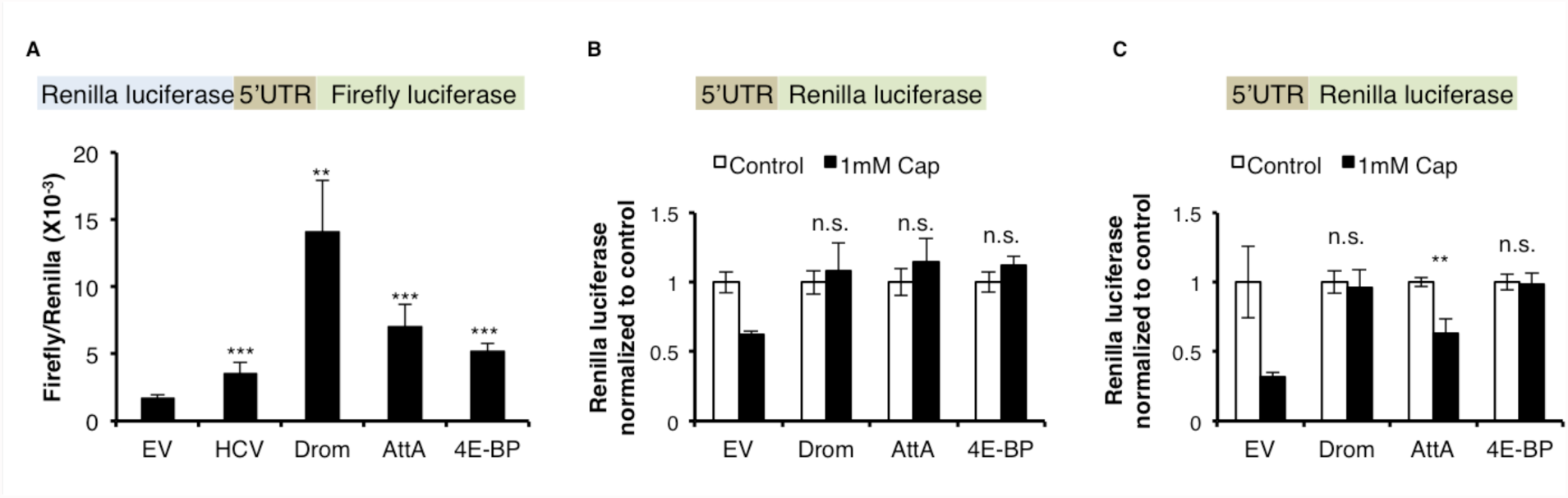
5’UTRs of Drosomycin, Attacin A and 4E-BP can be translated cap-independently. 3A. Schematic showing the biscistronic construct (top). (Bottom) Luciferase activity from S2 cells expressing the bicistronic construct the 5’UTRs from Drosomycin (Drom), Attacin A (AttA) and 4E-BP with the empty vector without any 5’UTR inserted (EV) as a negative control and Hepatitis C virus (HCV) IRES inserted as a positive control. Renilla luciferase activity of each sample is normalized to it’s respective Firefly luciferase activity. 3B. Monocistronic reporter (schematic on top) assays with indicated 5’UTRs on X-axis in control RRL (white) or RRL treated with excess cap-complex (black). Data for each 5’UTR are normalized to respective control samples. (EV= empty vector). 3C. Monocistronic assay as performed in 3B but in RRL treated with microccocal nuclease to eliminate competing capped transcripts. All data are mean of at least 3 independent experiments, error bars indicate standard error. All data are mean of at least 3 independent experiments, error bars indicate standard error. *=p<0.05, **=p<.01, ***=p<0.001. All p-values are calculated w.r.t. respective controls.

### Translational bias imposed by 4E-BP drives AMP synthesis

Data from Fig. 2 showing that AMP synthesis is reduced in 4E-BP mutants and Fig. 3 showing that AMPs are translated cap-independently together suggests a role for 4E-BP in biasing cellular translation to cap-independent mechanisms. To test the idea that 4E-BP is required for favoring cap-independent translation of AMPs during infection, we tested the bicistronic reporter (Fig. 3A) in S2 cells expressing a constitutively active phospho-mutant of 4E-BP, 4E-BP^LLAA^. The LLAA mutant cannot be phospho-regulated by the kinase mTOR, and thereby inhibits eIF4E more consistently when overexpressed [22]. The bicistronic reporter containing the 5’UTRs of Drosomycin, Attacin A and 4E-BP greatly enhanced the expression of the 2^nd^ cistron (reporting IRES) in the presence of 4E-B^PLLAA^ in comparison to cells expressing GFP as a control (Fig. 4A). We further corroborated this using a second bicistronic reporter with GFP reporting cap-dependent translation and dsRed reporting cap-independent translation (schematic, Fig 4B). S2 cells expressing the Drosomycin and Attacin A 5’UTR bicistronic reporters showed an enhanced dsRed expression when subjected to amino acid deprivation to activate GCN2. GFP levels remained unchanged, most likely due to the perdurance of GFP that was synthesized prior to the relatively short amino acid deprivation treatment (4 h). These data show that increasing 4E-BP levels in cells results in enhanced translation of transcripts containing cap-independent 5’UTRs.

**Fig. 4.**
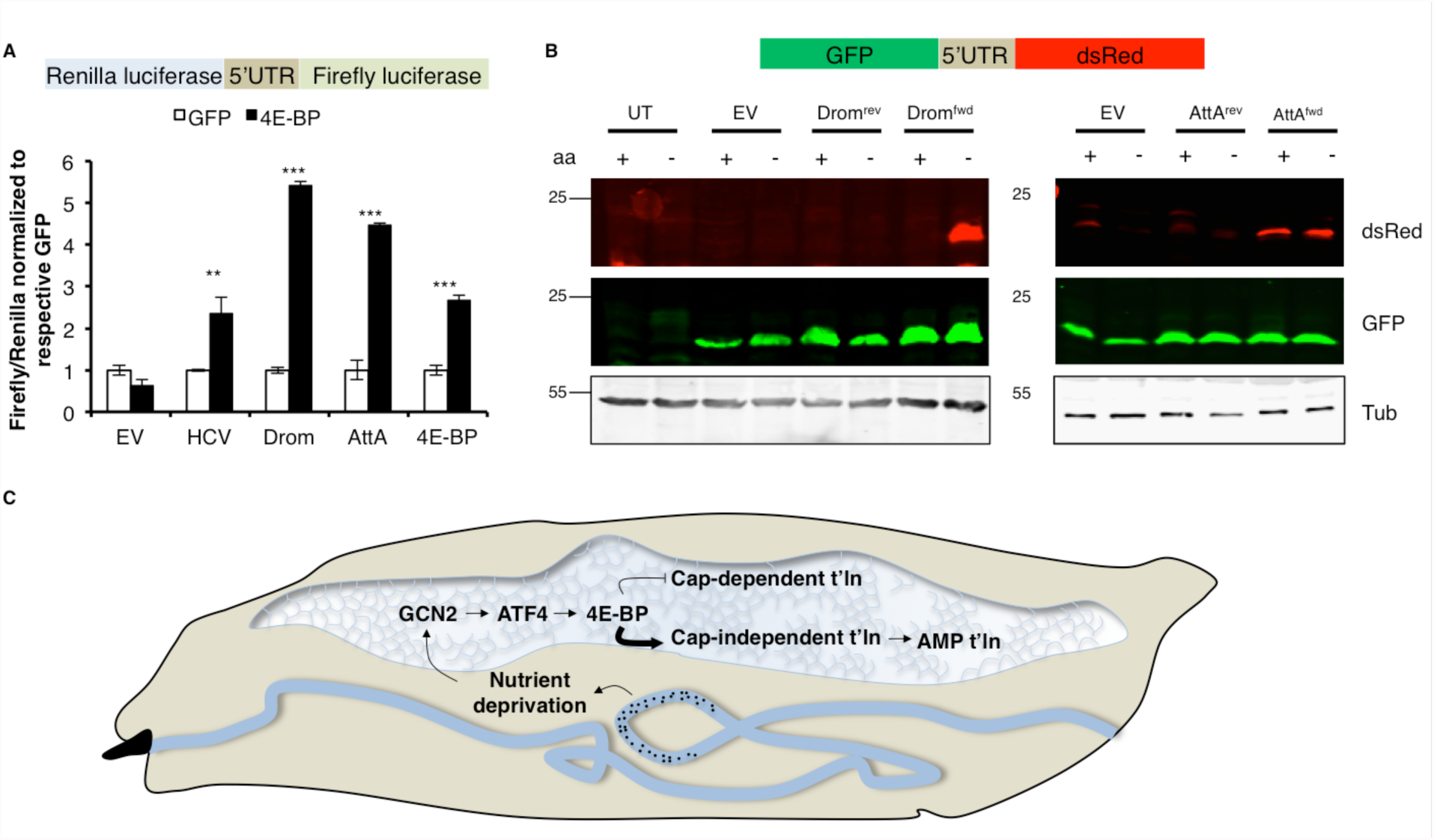
4E-BP enhances the translation of AMPs. 4A. Bicistronic reporter (schematic on top) assays with 5’UTRs indicated on X-axis in S2 cells overexpressing either GFP as a control (white) or 4E-B^PLLAA^ (black). Renilla luciferase activity of each sample is normalized to firefly luciferase activity of itself, and all samples are normalized to their respective control (GFP). 4B. Western blot analysis of the fluorescent bicistronic reporter (schematic on top) with Drosomycin (Drom) and Attacin A (AttA) 5’UTR in forward (^fwd^) or reverse (^rev^) orientation in S2 cells grown in either complete media (control) or media lacking amino acids (aa). (UT= untransfected control, EV= empty vector). 4C. **A model for 4E-BP activation and function during enteric infection in larvae.** In response to enteric infections by pathogens (indicated by black dots), GCN2 is activated in the fat body likely in response to nutrient deprivation induced by infection. GCN2 activation in the fat body leads to ATF4 synthesis and subsequent transcriptional induction of 4E-BP. Once induced, 4E-BP itself is synthesized cap-independently and blocks cap-dependent translation of most cellular transcripts. In doing so, it promotes (indicated by bold arrow) cap-independent translation of AMP transcripts. Such bias in cellular translation by 4E-BP is required to drive AMP synthesis during the infection, thus explaining the role of 4E-BP in the innate immune response. *=p<0.05, **=p<.01, ***=p<0.001. All p-values are calculated w.r.t. respective controls.

## Discussion

A cumulative body of research surrounds the transcriptional response to infection as mediated by the innate immune response pathways. Although translational inhibitors such as GCN2 and 4E-BP have been implicated in the antibacterial response, the incongruence of translation inhibition and the need for AMP synthesis had not been addressed. Here, we provide a basis for a new concept that AMPs have evolved mechanisms to bypass translation inhibition by 4E-BP, which is highly induced in hosts infected with pathogenic bacteria. This work also adds a new dimension to the ISR pathway by demonstrating its requirement in antibacterial response. Our data suggests that in the context of oral infections with a non-lethal pathogen, *Ecc15*, GCN2 is activated in the fat body (Fig 4C). Subsequent to GCN2 activation, ATF4 induces 4E-BP in the fat body, where it biases cellular translation towards cap-indepndent mechanisms that favor AMP synthesis (Fig 4C).

While there have been some clues about how enteric infections can be communicated to the fat body [23], which is known to be the primary site of AMP synthesis, the causes of GCN2 activation remain unclear. One possible explanation could be that infection leads to nutritional deprivation. This idea has been suggested before [7], but the downstream effects of such deprivation appear to vary depending on the infectious agent and the responding tissue. While infection of a severe pathogen such as *P.entomophila* results in a GCN2-mediated translation block in the gut that is detrimental to the host immune response [7], our observation is consistent with the positive effects of 4E-BP induction in the fat body in mounting an innate immunre response against a non-lethal pathogen such as *Ecc15* [5]. It is possible that the detrimental effects of GCN2 in the context of *P. entomophila* infection is mediated not through ATF4 and 4E-BP, but through other independent effectors.

It is worth noting that GCN2 engages two different translation inhibition mechanisms: 1) phospho-eIF2α, which induces ATF4 and, 2) 4E-BP, which is transcriptionally induced by ATF4 [12, 13]. ATF4 is a very effective transcription factor that induces expression of various stress-response genes including an eIF2α phosphatase subunit, GADD34 [24, 25], and a second translation inhibitor, 4E-BP. GADD34 itself is synthesized by an intriguing mechanism involving upstream uORFs (similar to ATF4) that are favorably translated under phospho-eIF2α conditions[26]. While GADD34 relieves the translation inhibition by phospho-eIF2α by feedback, inhibition by 4E-BP may persist longer. This is supported by our data showing that the 5’UTR of 4E-BP itself is synthesized favorably by cap-independent translation, suggesting that this inhibition mechanism is self-sustaining (Fig. 3, 4A).

In addition to regulation by GCN2/ATF4, 4E-BP is also famously regulated post-translationally by another amino-acid sensitive kinase, mTOR [27]. While under steady state conditions, mTOR phospho-inactivates 4E-BP, mTOR itself is inactivated in response to amino acid deprivation. Thus 4E-BP that is newly synthesized in response to infection mediated amino acid deprivation will likely not be subject to inactivation by mTOR.

Since suppression of cap-dependent transcripts appears to stimulate the synthesis of AMPs (Fig. 4A, B), we speculate that the 5’UTRs do not compete well with other cellular mRNAs for ribosomes and initiation factors in unstressed cells. According to this view, AMP translation may be significantly enhanced when 4E-BP inhibits translation of cap-dependent transcripts and the translation machinery becomes more available. In addition to the established role of 4E-BP in the inhibition of cap-dependent translation, our data indicates a more nuanced role for 4E-BP as a promoter of cap-independent translation required for driving the synthesis of essential immune response proteins. It is notable though that the 5’UTRs of Drosomycin and Attacin A are relatively small at 63 and 30 bases respectively. To the best of our knowledge, these are the smallest characterized cap-independent translation elements known. The most commonly studied cap-independent translation mechanism are IRES elements, which were first discovered in viral mRNAs, capable of recruiting 43S ribosomal subunits directly. While there are no known consensus sequences for IRESes, scoring positively in biochemical assays such as bicistronic assays and cap-competion assays has been recognized to be a reliable predictor for them [28]. In addition to IRESes, there are several other 5’UTR features that can promote cap-independent translation. For example, the presence of <u>C</u>ap-<u>i</u>ndependent <u>t</u>ranslational <u>e</u>nhancers (CITE) in 5’UTRs [29–31] or m6-adenylation of 5’UTRs[32, 33] have been shown to promote recruitment and assembly of translation initiation complexes. It is important to note that both these mechanisms require the 5’ end of the mRNA to be available, and thus transcripts bearing these features usually fail the bicistronic assay, indicating that at least the two AMPs (Drosomycin and Attacin) tested herein likely do not contain these mechanisms.

The eIF4F complex, of which eIF4E is a part, is probably the best studied cap-recognition complex. However, recent studies indicate that several transcripts are capable of being translated even when eIF4E is inactivated, due to cap-binding activity of another initiation factor, eIf3d [34]. Such transcripts are thought to contain eIF4F inhibitory elements that ensure that their translation occurs through a specialized pathway involving eIF3d, which is part of the larger eIF3 initiation complex. What comprises such inhibitory elements or whether there are other features in the mRNA that promote recruitment of eIF3 over eIF4F remain unknown. While our data indicates that Drosomycin and Attacin A are regulated by limiting eIF4E availability, the possibility that other AMPs are regulated by alternative cap-recognition mechanisms remains a possibility. It would be interesting to examine whether the mammalian innate immune response is aided by any of the three known mammalian 4E-BPs and if first-response cytokines are synthesized cap-independently.

## Methods

### Fly stocks and S2 cell culture

All *Drosophila* stocks were reared on standard cornmeal medium at room temperature. A list of stocks can be found in the Table S1.

S2 cells were grown on standard Schneider’s medium (Life Technologies) supplemented with 10% Fetal Bovine Serum and 1% penicillin/streptomycin. For transfection, cells were plated at 1.25x10^6^ cells/ml in Schneider’s supplemented with 10% FBS and an additional 1 μg/ml bovine insulin. DNA was transfected at a 5:1 ration of expression plasmid to reporter plasmid using polyethylene imine (PEI) or Effectene (Qiagen).

### Larval infection

Bacteria were grown in LB broth in an overnight culture and pelleted. All bacteria tested were grown at 37°C except *Ecc15* and *Serratia marcescens*, which were grown at 25°C. 4-day old larvae orally infected with food mixed 5:1 by weight with the bacterial pellet for 4 hours. A small amount of bromophenol blue dye was added to the food to enable selection of larvae that were successfully infected.

### Systemic pathogen load

After infection, the larvae were briefly washed in 70% ethanol to surface sterilize them. 3-4 larvae per condition were homogenized using a pestle in 100 μl of PBS. The homogenate was briefly centrifuged at 1000 rcf, 3 min to eliminate debris and serially diluted before plating on selective LB agar plates. For quantitation, colonies were counted from the same dilution for different conditions and genotypes and normalized to the control.

### Immunofluorescence (IF) and Western blotting (WB)

Larval guts and fat body were dissected in PBS and fixed in 4% PFA, 0.1% Tween, 1x PBS for 20 minutes. The tissues were then washed 3x in PBT (0.1% Tween) and stained with respective antibodies in dilutions indicated below. Tissues were mounted in 50% glycerol containing DAPI and imaged using a LSM700 Zeiss microscope at 20X magnification unless otherwise specified. Antibodies: mouse anti-GFP 1:500 for IF and 1:2000 for WB (A6455, Life Technologies), rabbit anti-dsRed 1:500 for IF and 1:2000 for WB (R10367, Thermo Fisher), mouse anti-actin 1:5000 WB (MAB1501, Millipore), mouse anti-attacin (Dr. Donggi Park) 1:500 WB.

### Molecular cloning

Luciferase bicistronic reporter: The 5’ UTR sequences for Drosomycin, Attacin A and 4E-BP, were subcloned into previously described luciferase bicistronic reporter constructs.

Fluorescent bicistronic reporter: The reporter backbone was generated in a pCASPER4 vector by sequentially cloning in a GFP (XbaI) and dsRed (EcoRI), with the NotI and SacII sites in between them available for insertion of various 5’UTRs. 5’UTRs of Drosomycin and Attacin A were assembled by extension PCR and inserted in the NotI/SacII sites in either the forward or reverse orientation.

### Hemolymph collection for WB and mass spectrometry (MS)

Infected larvae were bled into PBS by making a small incision and the diluted hemolymph was cleared of hemocytes by centrifuging at 1000g for 3 min at room temperature. 10-20 larvae for each condition were bled for WB and 75 larvae were bled for mass spectrometry. To prepare samples for MS, the diluted hemolymph was passed through a 10kDa filter (Millipore). The low mass filtrate was then reduced with dithiothreitol (2 μL of 0.2 M, pH 8) for 1 hr at 57°C and subsequently alkylated with iodoacetamide (2 μL of 0.5 M, pH 8) for 45 min in the dark at room temperature. Immediately following alkylation, peptides were desalted using a solid phase extraction (SPE) cartridge containing C18 resin. The resulting eluate was then concentrated to dryness and resuspended in 0.5% acetic acid for mass spectrometric analysis.

### Liquid chromatography and Mass spectrometry

Each sample was separated by reverse phase chromatography using an EASY-nanoLC 1000 system (Thermo Scientific) configured for preconcentration using an Acclaim PepMap trap column in line with an EASY-Spray 50 cm x 75 μm ID PepMap C18 analytical column (2 μm beads). Solvent A was 2% acetonitrile containing 0.5% acetic acid and solvent B was 80% acetonitrile with 0.5% acetic acid. Separation was carried out using a linear gradient from 5-35% solvent B over the first 60 min and then increased to 45% B over 10 min. The gradient was then ramped to 100% B over 10 min and held constant for an additional 10 min. The LC system was coupled via an EASY nano-electrospray ionization source (Thermo Scientific) maintained at 2.0kV to a Q-Exactive Orbitrap mass spectrometer (Thermo Scientific). For all experiments, high-resolution MS1 spectra were acquired with a resolving power of 70K (at m/z 200), automatic gain control (AGC) target of 1e6, maximum ion time of 120 ms, and scan range of 400 to 1500 m/z. Following each MS1 scan, data-dependent acquisition of high resolution HCD MS2spectra were acquired for the top 20 most abundant precursor ions in the preceding full scan. All MS2 spectra were collected using a single microscan at 17,500 (at m/z 200), AGC target of 5e4, maximum injection time of 120 ms, 2 m/z isolation window, and Normalized Collision Energy (NCE) of 27. All LC-MS/MS data was searched using a standalone version of the Byonic algorithm (Protein Metrics Inc.) against a *Drosophila* UniProt database modified to include known antimicrobial and immune-induced peptide/protein sequences. All data was analyzed using a no enzyme (non-specific) search with a precursor mass tolerance of ±10 ppm and fragment ion mass tolerance of ±10 ppm. Carbamidomethylation of Cys was added as a static modification and oxidation of methionine and deamidation of glutamine and asparagine were searched as a variable modification. The results were filtered to only include peptides identified with a Byonic score of 300 or better.

### Bicistronic assays

36 hours after transfection with the luciferase bicistronic reporter, cells were lysed in passive lysis buffer (Promega) and luciferase activity was measured by a dual luciferase assay. Firefly expression was measured in 75 mM HEPES pH 8.0, 5 mM MgSO4, 20 mM DTT, 100 μM EDTA, 530 μM ATP, 0.5 mM coenzyme A, and 0.5 mM D-luciferin. Renilla expression was measured by addition of an equal volume of 25 mM Na4PPi, 10 mM NaOAc, 15 mM EDTA, 0.5 M Na2SO4, 1.0 M NaCl, and 0.1 mM Coelenterazine.

For experiments in Fig. 4A, cells were co-transfected with either pMT-GFP or pMT-4E-BP^LLAA^ and expression of GFP or 4E-BP was induced with 0.5mM copper sulfate for 2 h after which the cells were harvested for luciferase assays as described above.

For experiments in Fig 4B, cells transfected with the fluorescent bicistronic reporter were harvested in lysis buffer containing 1% NP40, 1x TBS and protease inhibitor cocktail (Roche) and subjected to western blotting.

### In vitro Transcription and mRNA reporter preparation

Transcription templates for monocistronic reporters were created using PCR with a template specific forward primer containing a T7 promoter sequence and a vector specific reverse primer. Templates were transcribed using T7 RNA polymerase and purified using LiCl precipitation. Transcripts were capped using Vaccinia Virus Capping enzyme (New England Biolabs, Ipswitch, MA) as recommended by the manufacturer and purified by phenol/chloroform extraction and isopropanol precipitation. Transcripts were poly(A) tailed using E. coli poly(A)polymerase (New England Biolabs, Ipswitch, MA) and purified by phenol/chloroform extraction and isopropanol precipitation.

### In vitro translation assays

Translation assays were performed in 10 μl reactions containing: 6 μl of rabbit reticulocyte Lysate (Green Hectares, McFarland, WI) (either treated with Micrococcal nuclease to eliminate endogenous mRNAs or untreated to allow for translation under competitive conditions), 0.1 mM spermidine, 60 μm Amino Acids, 16.8 mM creatine phosphate, 800 ng of creatine kinase, 24 mM HEPES (pH 7.4), 0.4 mM Mg acetate, 30 mM K acetate, 1 μg of calf liver tRNA, 2 units SUPERase-In RNase Inhibitor (ThermoFisher, #AM2696), and 100 ng of template RNA. Translation reactions were incubated at 37°C for 30 min and luciferase activity was measured using 100 μl of luciferase substrate (Promega). In experiments containing excess m7G cap, a cap structure analogue (New England Biolabs, #S1407S) was added to a final concentration of 1 mM. All experiments were performed at least twice in triplicate.

## Acknowledgements

We thank the NYU Medical Center Proteomics Core, particularly Victoria Cotham (for her help with mass spectrometry analyses), Mimi-Shirasu Hiza and Brian Lazzaro (for providing the pathogens used in this project), Neil Silverman and Donggi Park (for providing the anti-Attacin antibody), Marc Amoyel, Erika Bach and Jessica Treisman (for discussions and advice), and the Bloomington Drosophila Stock Center and the Vienna Drosophila Stock Center for fly lines used in this project. This work was supported by grants from the National Institutes of Health (R01EY020866) and the March of Dimes (FY#13-204) to H.D.R., the National Institutes of Health (R01GM11703) and a New Scholar in Aging Award from the Ellison Medical Foundation to M.T.M., and the American Heart Association fellowship (17POST33420032) to D.V. The mass spectrometric experiments were supported in part by NYU School of Medicine.

## Author contributions

D.V. and H.D.R. conceived the project, designed the experiments, analyzed the data, and wrote the paper. N.C. and M.T.M. performed the in-vitro IRES assays. J.S. provided technical assistance for all fly experiments. B.U. generated the mass spectrometry data.

